# Expectation effects on repetition suppression in nociception

**DOI:** 10.1101/2025.07.11.664149

**Authors:** Lisa-Marie G. Pohle, Moritz M. Nickel, Birgit Nierula, Markus Ploner, Ulrike Horn, Falk Eippert

## Abstract

Repetition suppression, the reduced neural response upon repeated presentation of a stimulus, can be explained by models focussing on bottom-up (i.e. adaptation) or top-down (i.e. expectation) mechanisms. Predictive coding models fall into the latter category and propose that repetitions are expected and therefore elicit smaller prediction error responses. While studies in the visual and auditory domain provide some support for such models, in nociception evidence remains inconclusive, despite the substantial influence expectations exert on pain perception. To assess expectation effects on repetition suppression in nociception, we developed a paradigm in which healthy volunteers received brief CO_2_ laser stimuli, while we acquired electroencephalographic (EEG) and peripheral physiological data. Importantly, laser stimuli could be either repeated after one second or not be repeated, with the probability of repetitions manipulated in a block-wise fashion, such that repetitions were either expected or unexpected. We observed repetition suppression in laser-evoked potentials as well as laser-induced gamma band oscillations, but not in laser-induced desynchronisations in the alpha and beta band. Critically, neither these EEG responses, nor the peripheral physiological data showed significant differences between the expectation conditions, with Bayesian analyses mostly providing evidence for an absence of effects. This indicates that repetition suppression to brief nociceptive laser stimuli is not driven by top-down factors, but rather mediated by other adaptation processes. While this does not preclude an influence of predictive coding models in nociception, it suggests that when the nervous system receives highly precise input, its responses are less susceptible to influence from expectations.

## INTRODUCTION

Repetition suppression, the reduced neural response to the repeated presentation of a stimulus, is a fundamental phenomenon of sensory processing (1). While consistently observed across sensory domains (for evidence in the olfactory, somatosensory, visual, and auditory domain, see e.g., 2–5), the underlying physiological mechanisms remain a subject of debate (6–8). Suggested mechanisms can be broadly categorised into two branches: on the one hand, bottom-up approaches like receptor habituation or neuronal fatigue, as well as sharpening or facilitation, are able to explain repetition suppression (1). On the other hand, top-down approaches such as predictive coding argue that the first occurrence of a stimulus happens at an unpredictable time point, leading to greater surprise or prediction errors than the repetitions, where stimulus occurrence, characteristics and timing are more predictable (9).

One approach to distinguish between those explanatory frameworks is to vary the probability of a repetition, i.e. p(rep). If repetition suppression is mainly driven by bottom-up processes, this variation should not exert any influence. If repetition suppression is however driven by top-down factors such as expectation, different p(rep) should lead to different magnitudes of repetition suppression (e.g. larger p(rep) should lead to larger repetition suppression). Evidence for a modulation of repetition suppression by p(rep) can indeed be observed in the visual and auditory domain, but this strongly depends on contextualizing factors such as specifics of the employed paradigm and stimuli (e.g., 5,10–17).

In the nociceptive domain, expectation effects on repetition suppression have rarely been studied, despite the strong impact expectations exert on pain (18). Researchers have utilized the high temporal precision of nociceptive laser stimuli, which yield well characterised electroencephalography (EEG) responses. These include the lateralized N1 component, the later N2P2 complex or vertex wave (19,20), as well as induced desynchronizations in the alpha and beta band and synchronisation in the gamma band (21). Studies have found mixed evidence for a modulation of repetition suppression by expectations: while repetition suppression to laser stimuli is influenced by *temporal* expectations (22), expectations regarding other stimulus properties, such as *location* or *modality*, did not significantly influence repetition suppression (23,24).

Critically though, three important aspects of an expectational modulation of repetition suppression have not been assessed in nociception so far. First, the most fundamental manipulation, i.e. varying the probability of the repetition itself – i.e. p(rep) and thus the expectation – has not been carried out. Second, the above-mentioned studies did not investigate responses in the gamma frequency range, which are a core feature both of pain perception (25–28) and predictive coding (e.g., 21,29–31), where they have been associated with bottom-up (and thus in the predictive coding framework prediction error) signalling. Third, the absence of expectation effects on repetition suppression has not been formally assessed, as the studies above relied on frequentist statistics and where thus not able to provide evidence of absence, as possible via Bayesian statistics.

Considering that repetition suppression is a fundamental feature of sensory processing, we here set out to address these three open aspects in order to gain a deeper insight into the mechanism that underlie repetition suppression in the nociceptive domain. Specifically, we investigated repetition suppression in two different contexts, in which either the presentation of a *stimulus pair* (i.e. a repetition) or of a *single stimulus* (i.e. an omission of the second stimulus) was more likely. We expected that brief laser stimuli would elicit EEG responses, that should be weaker in response to the second stimulus, thus demonstrating repetition suppression. Furthermore, if nociceptive repetition suppression is driven by expectation, we would expect repetition suppression (i.e. the reduction in response to second stimulus) to be stronger in contexts with high p(rep).

## METHODS

### Participants

We invited 58 healthy volunteers (33 female, mean age: 26.3 years, age range: 18-35 years) who were recruited via the participant database of the Max Planck Institute for Human Cognitive and Brain Sciences. Participants did neither have a history of neurological or psychiatric disorders, nor acute or chronic pain. All participants provided written informed consent, and the study was approved by the Ethics Committee at the Medical Faculty of the Leipzig University.

No EEG data were acquired from 15 participants, as their responses in a pre-experimental stimulus calibration phase prevented them from participating in the main experiment: they either reported the minimally required stimulus temperature of 53°C as too painful (pilot data had shown that a temperature of 53°C was necessary for eliciting robust Aδ LEPs with our stimulation and recording set-up) or reported the maximum stimulus temperature of 59°C as not painful (a painful percept was deemed necessary to obtain robust responses in the gramma frequency range (25)). A further seven participants were excluded after data acquisition, as three participants did not reach minimum task performance (described below), three participants did not finish the experiment, and one participant had insufficient data quality (i.e. less than half of the trials remained after preprocessing). The final sample size thus consisted of 36 participants with complete datasets (17 female, mean age: 26.7 years, age range: 18-35 years), which would allow us to detect a medium-sized effect (d = 0.5) in a two-tailed paired-samples t-test with 90% power at an alpha-level of 0.05.

### Stimuli

Painful heat stimuli were delivered to a 5×5 cm area on the dorsum of the left hand via a CO_2_ laser with a wavelength of 10.6 µm (Laser Stimulation Device, SIFEC s.a., Ferrières, Belgium). The beam diameter was set to 6 mm and the stimulus duration to 125 ms. The device utilizes a closed loop temperature control system to maintain constant skin temperature during stimulation by adjusting the energy output. The stimulus position was set by an electric motor which moved the laser head relative to a participant’s hand and allowed for precise control of stimulation position. Participants’ vision was shielded, so that they were not aware of the exact stimulus location. Throughout the entire experiment, participants wore protective goggles.

The temperature used for each participant was determined in a pre-experimental two-step calibration procedure. During calibration, only repeated stimuli were used: first, one stimulus was applied at a random position within the stimulation area, which was then followed by an inter-stimulus interval of 1 s in which the laser beam was shifted 1 cm distal and a second stimulus with the same characteristics applied at the new position. After each trial (encompassing two stimuli), average ratings were acquired on a visual analogue scale (VAS) anchored at ‘No percept’ (left, converted to 0), ‘Pain threshold’ (middle, converted to 50) and ‘Unbearable pain’ (right, converted to 100).

In the first part of the calibration procedure, the temperature was increased for each trial (from a minimum of 40°C to a maximum of 59°C, with steps of 2°C for temperatures below 50°C and increments of 1°C for temperatures above) and participants rated their sensation. This part ended when a trial was rated as 90 or higher or the maximum temperature was reached. A linear regression was then fitted to determine the temperature reflecting the approximate pain threshold and the temperature leading to a rating of 90.

In the second part of the calibration procedure, those two temperatures (limited by a maximum of 59°C due to safety reasons), were then used as the extrema for the following stimulation (except when the distance between the pain threshold and 59°C would be smaller than 4°C, which would lead to 55°C being used as the minimum temperature to allow for the range of tested temperatures to be broad enough for the subsequent regression). The space in between those two temperatures was then divided evenly to end up with ten different temperatures, each of which was applied twice on different skin spots to account for spatial variability in sensitivity (with the order of applied temperatures shuffled). After acquiring those 20 trials, another linear regression was fitted to determine the temperature equivalent to a rating of 65, which was the aimed-for target in the main experiment (again limited by a maximum temperature of 59°C).

### Experimental procedure

The experiment started with the aforementioned calibration. This was followed by a longer break for the preparation of the data acquisition setup. The main experiment consisted of one block of a control task, followed by eight blocks of the main task and by another block of the same control task; each block lasted 4-5 minutes (Fig. 1a).

**Figure 1.**
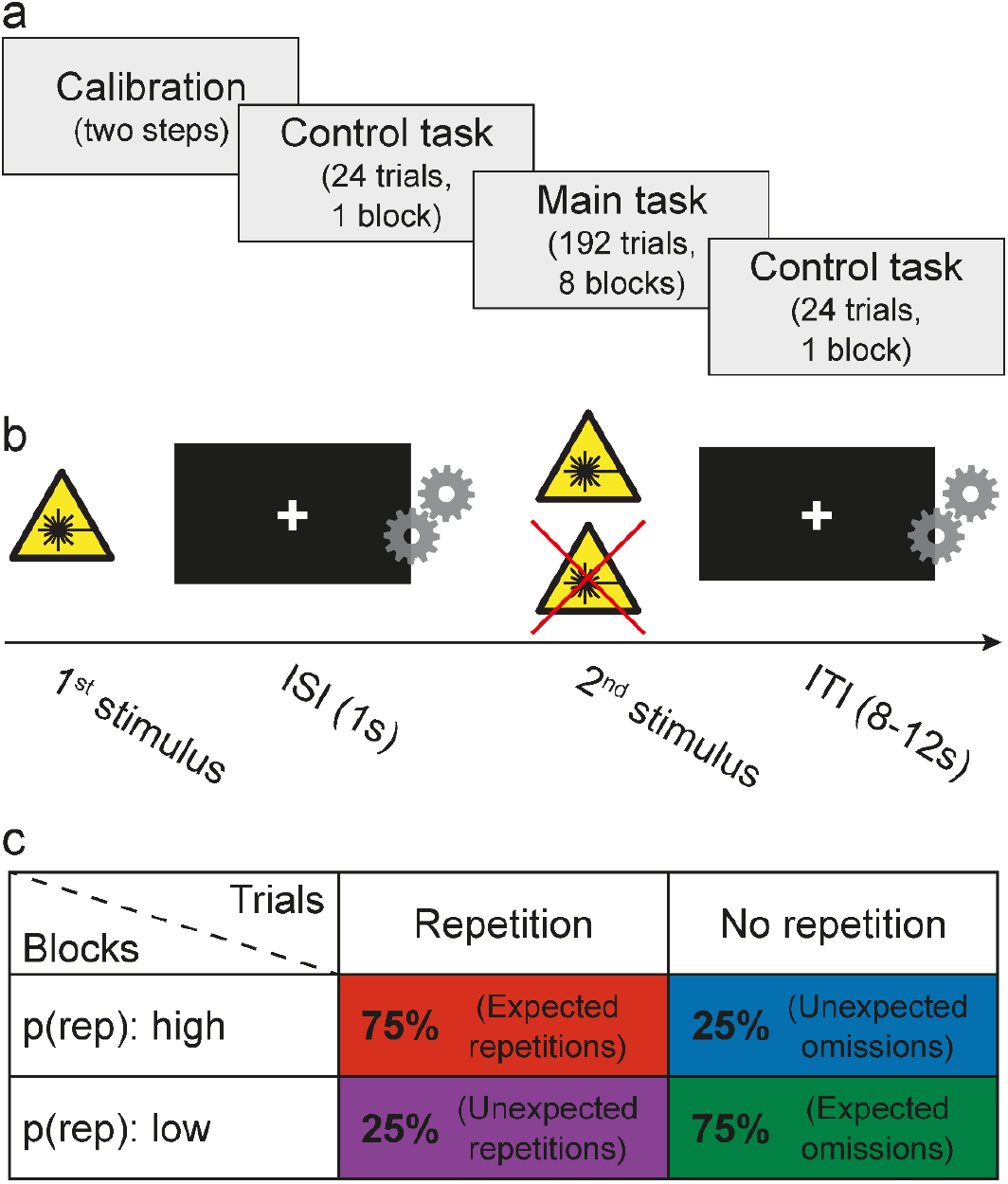
(a) General timeline of experiment. (b) Trial structure of main experiment. Each trial started with the application of a 125ms stimulus followed by an inter-stimulus interval of 1s, after which the stimulus could either be repeated or not repeated. During the inter-trial interval (duration randomly spaced between 8 and 12s) the laser was shifted to a new position. (c) Contingencies in the main experiment.

The main task trials could either be repetitions as described earlier (i.e. two stimuli being applied per trial), or omissions (i.e. only one stimulus being applied per trial). Both trial types could occur under two different conditions: either in a block, where repetitions were more likely and therefore expected (while omissions were rare and therefore unexpected; termed “high block”) or in a block, where repetitions were rare and therefore unexpected (while omissions were more likely and therefore expected; termed “low block”). Each block consisted of 24 trials and every trial started with the delivery of one laser stimulus, which was either repeated after 1s (repetition trial) or was not repeated (omission trial). During this ISI, the laser beam was shifted 1cm anterior irrespective of the occurrence of a repetition to avoid any influence of the sound emitted by the electric motors. Trials were separated by an interval chosen randomly from the range 8-12s (uniform distribution) during which the laser beam was shifted to a new random position (Fig. 1b). We varied the repetition probability between blocks, with high blocks consisting of 75% repetitions and 25% omissions and vice versa for low blocks (Fig. 1c). Before the beginning of each block, we informed participants about the type of the upcoming block, without disclosing the exact repetition probability (i.e. “In the following block repetitions will occur frequently/rarely”). The order of trials within each block was pseudorandom, with the constraint of always having at least three standard (or expected) trials between unexpected repetitions/omissions. The order of blocks was always alternating between the two types with the condition of the first block being balanced across participants. In total, this resulted in 72 expected repetitions/omissions and 24 unexpected repetitions/omissions across the experiment (the actual numbers could be lower on an individual level due to technical difficulties, mainly laser aborts in case the distance to the target was beyond the operational range).

To assess whether participants formed the intended expectations, before the start of each block of the main task we asked them to rate how often they expected to experience repeated stimuli in the upcoming block. We used a VAS anchored at ‘Never’ (left, converted to 0%) and ‘Always’ (right, converted to 100%). Furthermore, at the end of each block we asked participants to rate their average pain perception across the block (using the earlier-described VAS for pain ratings) - crucially, they had to provide two separate ratings for the first stimulus of each trial and the repetitions.

In the control task we aimed to assess whether participants were able to distinguish between repetitions and omissions (participants who were not able to do so were excluded from the experimental sample; N=3, see also section ‘Participants’). Therefore, we used trials with the same structure as in the main experiment but set the repetition probability to 0.5. After each trial and a pause of 1.5s participants had to indicate within a time-window of 2s whether they perceived one stimulus or two stimuli.

### Recordings

A 32-channel electroencephalogram (EEG) was recorded using a Bittium NeurOne EEG system (Bittium Corporation, Oulu, Finland). We used a cap with passive Ag/AgCl electrodes positioned according to the international 10-20 system and referenced to a nose electrode (BC-TMS-32, Easycap, Wörthsee, Germany). To allow for correction of ocular artifacts, we further recorded an electrooculogram (EOG) using two additional electrodes placed on the outer canthus of the right eye (referenced to nose) and below the right eye (referenced to Fp2). We used a sampling rate of 2kHz with a hardware anti-aliasing filter at a cut-off frequency of 500 Hz and all recordings were grounded at POz, with impedances aimed to be below 5 kΩ.

In addition to the EEG data, we also acquired peripheral physiological data (pupil diameter and skin conductance responses; cardiac and respiratory data were also acquired, but not analysed here), as these allowed us to assess the effects of expectations on a further processing level. Pupil diameter was recorded from the left eye using an EyeLink 1000 system (SR Research, Ottawa, ON, Canada) with a sampling rate of 1 kHz with a hardware low-pass filter and a cut-off frequency of 250Hz; participants rested their head on a table-mounted chinrest. Skin conductance was recorded with a BrainAmp ExG system (BrainProducts, Gilching, Germany) at a sampling rate of 1 kHz using Ag/AgCl electrodes filled with isotonic electrolyte gel and attached to the thenar and hypothenar eminences of the left hand.

### EEG data processing

All processing and analysis code is available at https://github.com/eippertlab/eeg-repetition-suppression. EEG data were mainly processed using MNE-Python (version 1.2.2, 32), supplemented by FieldTrip (version 0.20220623, 33) for time-frequency decomposition and analysis. Data from the eight experimental blocks were concatenated and downsampled to 500Hz (anti-aliasing filter at a cut-off frequency of 250Hz).

For the analysis of LEPs, data were band-pass filtered (1-30Hz, using a 4^th^ order Butterworth filter) to remove slow drifts and suppress muscle artifacts. Following this, data were epoched in a window between 2s before and 4s after the onset of the first stimulus of each trial. Invalid trials (e.g. aborts of the laser) were removed and the remaining epochs were cleaned from eye blinks and movements via independent component analysis (ICA) using the extended infomax algorithm (34). Finally, data were re-referenced to the average of all channels for the analysis of the N2P2 component and to Fz for the analysis of the N1 component and baseline corrected relative to the 500ms before stimulus onset. Trial counts of each condition were equalized per participant by selecting the trials with the greatest temporal proximity to the trials of the condition with the lowest trial count, resulting on average in 21.1 trials per condition per participant (range 13-24). Average waveforms were calculated per condition and participant. Furthermore, amplitudes of the N1 (at T8-Fz), N2 and P2 (both at Cz-average) components were extracted per participant and condition via the following procedure: since the P2 was found to be the most prominent component, we first determined the latency of the highest deflection during an interval of 100ms around the group level P2 latency. Then, we searched for the lowest deflection in the 200ms preceding the individual P2 to get the N2 latency. Amplitudes were then extracted by averaging the data in a 30ms window around this peak for the first stimulus and in the same time window shifted by 1 s for the second stimulus (or possible omission response). For the N1, we searched for the lowest amplitude during the 100ms before the N2 of each participant and extracted the average of the 30ms around this peak.

For the time-frequency analysis, a more extensive data cleaning procedure was implemented to thoroughly address myogenic noise (likely caused by the seating position necessary for eyetracking) as well as environmental noise (arising from the use of passive electrodes in a non-shielded environment). Strict low-pass filtering as carried out for LEPs was not possible, since we were also interested in oscillations in the gamma frequency range (30-100 Hz). Therefore, data were only high-pass filtered at 1 Hz (using a 4^th^ order Butterworth filter) and notch filtered around 50Hz and harmonics (using an 8^th^ order Butterworth filter). Subsequently, data were epoched and selected as described above, followed by a manual removal of strongly noise contaminated epochs and channels as determined by condition-blind visual inspection. The epoched data were then subjected to ICA (using the extended infomax algorithm, 34) and a recently-developed spike detection tool was used within IC-space to remove muscle artifacts: this is based on i) searching for locally extreme spikes in the high-pass filtered data within a sliding window and ii) removing them by fitting a Gabor function to this spike and subtracting it from the original data (35). Next, ICs associated with ocular or muscle artifacts were removed from the data. After ICA, epochs presenting with amplitudes exceeding ±100µV as well as jumps between adjacent datapoints exceeding 50µV were automatically discarded. Epochs were then passed to FieldTrip for time-frequency decomposition (see below) and visually inspected (by two independent and condition-blind raters) on a single-trial basis to detect remaining muscle artifacts and exclude the contaminated epochs. Trial counts of each condition were equalized per participant using a method of greatest temporal proximity, resulting in 18.4 trials per condition per participant on average (range 11-23).

To calculate a time-frequency representation (TFR) of the data, we used a fast Fourier transform on the Hanning-tapered data with a window length of 500ms for frequencies <=30Hz and a window length of 250ms for frequencies between 30-100 Hz. Those single-trial representations were then baseline corrected via subtraction of the average power in the period from 750ms to 250 ms before the first stimulus; a subtractive rather than a divisive approach was used to prevent an artificial increase of synchronisations (36). For further analysis, average values in the following temporal-spectral regions of interest (ROIs) were extracted for each participant and condition: alpha (500-1000 ms, 8-12 Hz), beta (300-700 ms, 13-30 Hz) and gamma (150-450 ms, 70-90 Hz). The choice of time windows and frequency bins followed Nickel et al. (37), with the only addition being that we prolonged all windows by 100ms to account for our longer stimulus duration.

### Peripheral physiological data processing

#### Skin conductance

Skin conductance data were processed with in-house Python scripts utilizing MNE and SciPy functionality. Data were filtered with a bidirectional first-order Butterworth bandpass filter and cut-off frequencies of 5 Hz and 0.0159 - corresponding to a time constant of 10s - and down-sampled to 10Hz (38). Data were cut into epochs ranging from −1s to 8s relative to the first laser stimulus of each trial and were baseline-corrected (at trial-time 0ms) for visualization purposes only. Considering that it is a common occurrence for participants to not show skin conductance responses, we distinguished between responders and non-responders by assessing whether a participant had at least one run per condition that contained responses to at least 25% of the trials. The minimum amplitude change necessary for a response was defined as an amplitude increase of 0.01 μS in a response window of 1 to 8s after the first laser stimulus in comparison to an average baseline amplitude in a baseline window −1 to 0s before the first laser stimulus (39,40). If participants did not reach these thresholds they were removed from the analysis as non-responders (5 of 36, leaving N=31 for SCR analyses). Trial counts of each condition were again equalized per participant using the aforementioned method of greatest temporal proximity before creation of the grand average plots. Trial-wise peak amplitudes were extracted by searching for a maximum within an interval of 1s to 6s after the first laser stimulation and subtracting the minimum that can be found within an interval from 0s up to this maximum. For further statistical analysis, these amplitudes were averaged within each condition.

#### Pupil dilation

Since eye tracking data could not be recorded in two participants, the pupil dilation analysis encompassed 34 participants. Eye tracking data were initially processed using MNE-Python to linearly interpolate missing data due to blinks as detected by the EyeLink Software in an interval from 50ms before to 200ms after the blink event. Subsequently, pupil data were processed with in-house Python scripts utilizing MNE and SciPy (41) functionality. The data were first filtered with a first-order low-pass Butterworth filter with a cut-off frequency of 2 Hz, and remaining blink artifacts were manually detected. The rater was blind to event timing and trial types during this process. Data were cut into epochs ranging from −1s to 8s relative to the first laser stimulus of each trial and were baseline-corrected (at trial-time 0ms, i.e. the onset of the first laser stimulus) for visualization purposes only. Epochs that contained more than 50% interpolated data points were excluded from the analysis (3.7% of all data). To account for differences in measured pupil dilation due to factors like brightness in the room, we z-standardized all values in the epochs within each participant before creation of grand average plots. Trial counts of each condition were equalized per participant using the aforementioned method of greatest temporal proximity. Trial-wise peak amplitudes were extracted by searching for a maximum within an interval of 1s to 6s after the first laser stimulation and subtracting the minimum that can be found within an interval from 0s up to this maximum. For further statistical analysis, these amplitudes were averaged within each condition.

### Preparatory data analysis

#### Correct response rates

The percentages of correct responses in the control task blocks were calculated for each participant and condition (i.e. one stimulus or two stimuli). Trials in which the laser aborted were excluded. If no response was given in the prespecified time window this was counted as a wrong response. Participants (N=3) achieving less than 50% correct answers in either of the conditions across both blocks were excluded from further analysis and are not contained with the 36 participants upon which this manuscript is based.

#### Pain ratings

We assessed whether the participant-wise average ratings on the VAS were higher than the pain threshold (rating of 50) using a one-sample t-test. Additionally, average ratings per participant, block type (high vs. low, i.e. more or less overall stimuli) and stimulus position (first stimulus vs. second stimulus) were calculated and the subjected to a two-factor repeated-measures analysis of variance (ANOVA). Note that more sophisticated analyses of e.g. expectation effects were not possible due to the acquisition of pain ratings only at the end of each block instead of trial-wise (a choice motivated by the necessity to not interfere with the recording of autonomous nervous system responses).

#### Expectancy ratings

Ratings on the VAS were averaged per participant and block type (high vs. low) and then assessed to be above or below the chance level of 50% respectively using one-sample t-tests.

#### EEG data habituation

Visual data inspection revealed a substantial habituation of the EEG responses, which we aimed to statistically assess by comparing the first half of the main task (blocks 1-4) to the second half (blocks 5-8) using one-sided paired samples t-tests or Wilcoxon signed-rank tests (in case of non-normal data). We decided on splitting the experiment in two parts instead of more parts to keep trial counts in the smallest conditions as high as possible. For this comparison, we extracted the amplitudes and power values as described above once for the first four blocks and once for the last four blocks. These analyses were based on responses to the first stimulus (thus each trial per participant was used instead of equalizing the trial counts between the conditions) and revealed a reduction of the N1 amplitude by 45.9% (t_(35)_ = 3.2, p = 0.001), of the N2P2 amplitude by 39.8% (W = 660, p < 0.001), of the alpha desynchronization by 74.6% (W = 209, p = 0.026), of the beta desynchronization by 25.3% (W = 204, p = 0.021), and of the gamma band oscillations by 67.5% (W = 113, p < 0.001). Therefore, we decided to base all further analysis only on the data from the first half of the experiment (number of trials per participant and condition for LEPs: mean = 10.8, range = 6-12; for TFA: mean = 9.2, range = 3-12). Results using the full dataset can be found in the Supplementary Material and are largely similar to the data reported in the main manuscript. Note that splitting the data into e.g. thirds or quarters was not considered as the stimulus numbers would have been too low for robust statistical analyses.

#### Peripheral physiological data habituation

In accordance with the EEG data, we observed substantial habituation in the pupil dilation and skin conductance responses and aimed to quantify this by extracting amplitudes for the first and second half of the experiment. As these peripheral physiological responses are unfolding on a much slower timescale than EEG responses, we could not distinguish responses to first and second laser stimuli and instead could only observe combined responses when two stimuli were present in a trial. To nevertheless quantitatively assess habituation, we therefore performed a two-factorial repeated-measures ANOVA on the amplitudes for first and second half of the experiment for repeated and single stimuli separately (i.e. using two factors: trial type (repeated trial vs. single trial) and experimental half (1^st^ half vs. 2^nd^ half)). The effect of trial type on the amplitude was significant (pupil dilation: F(1,33) = 67.725, p < 0.001, η^2^ = 0.324; skin conductance: F(1,30) = 17.101, p < 0.001, η^2^ = 0.101), as – similar to the EEG data – was the effect of experiment half, thus demonstrating substantial habituation (pupil dilation: F(1,33) = 16.714, p < 0.001, η^2^ = 0.134; skin conductance: F(1,30) = 16.641, p < 0.001, η^2^ = 0.235); no interaction between the two factors was observed. In line with the EEG data, we therefore base all further analysis of the peripheral physiological data only on the data from the first half of the experiment. Results using the full dataset can be found in the Supplementary Material and are largely similar to the data reported in the main manuscript.

### EEG data analysis

Repetition suppression was quantified descriptively by calculating the percentage of response reduction when comparing responses to the first and the second stimulus in repetition trials. This was carried out for N1 and N2P2 (peak-to-peak amplitude), as well as for alpha, beta and gamma power. For statistical analysis, we compared responses to the first stimulus and to the repetition by means of paired-sample t-tests with the directed hypothesis, that the response to the second stimulus is smaller (for positive signed responses: N2P2, GBOs) or greater (for negative signed responses: N1, alpha and beta desynchronization) than the response to the first one.

To investigate influences of expectations on repetition suppression, we first quantified repetition suppression by either calculating the difference in the response amplitude between first and second stimulus (for responses with a positive amplitude: N2P2, GBOs) or the difference between second and first stimulus (for responses with a negative amplitude: N1, alpha and beta desynchronization). Using paired-sample t-tests, those differences were then compared between the high and low blocks, translating to expected and unexpected repetitions, respectively. We tested one-sided with the alternative hypothesis being that the repetition suppression is stronger in the expected than in the unexpected condition for all outcome measures.

To furthermore assess the presence of omission responses, which would only be expected to exist if a prediction about an upcoming stimulus is violated (i.e. only for unexpected omissions of the second stimulus), we compared the responses to omissions between the expected and unexpected condition (low vs. high blocks) using paired-samples t-tests (in both time- and time-frequency domain). As this is not the main focus of the study, results can be found in the Supplementary Material.

To be sensitive to effects outside our predefined ROIs, cluster-based permutation tests (42) on the whole time-domain waveforms as well as TFRs were carried out for the expected vs. unexpected contrast for repetition and omission trials separately. In the time-domain, we searched for temporo-spatial clusters using the data with average reference, while we searched for three-dimensional clusters (time, frequency, channel) in the TFRs. For each time-channel or time-frequency-channel value a single test statistic is calculated and clusters of adjacent datapoints with significant values of the test statistic are created. This procedure is repeated 10000 times with labels (expected vs. unexpected) being assigned randomly to the data. Using the distribution resulting from those permutations as a null distribution, significance of the found clusters can be established.

### Peripheral physiological data analysis

As alluded to in the previous paragraph, it is not possible to investigate the amount of repetition suppression in the peripheral physiological data as the responses to the laser stimuli are too close in time and overlap. It is however possible to test whether expected and unexpected repetitions differ by looking at the difference in the combined response between those trial types (after matching trial numbers of conditions; similar analyses for omission responses can be found in the Supplementary Material). This was carried out via one-sided paired-sample t-tests with the alternative hypothesis being, that the response is higher in the unexpected than in the expected condition: in repetition trials, repetition suppression should be stronger in the repetition-expected condition (high blocks; leading to smaller overall responses compared to low blocks), whereas in omission trials, omission responses should be stronger in the omission-unexpected condition (high blocks; leading to larger overall responses compared to low blocks).

To be sensitive to effects aside those affecting the maximum amplitude, cluster-based permutation tests (42) on the whole average time courses similar to the EEG analyses were carried out for the repeated vs. single stimuli and for the expected vs. unexpected condition (for repetition and omission trials separately). We searched within a time window of 1s to 8s after the first laser stimulation for clusters of data points that have a significant value of the test statistic. To correct for multiple comparisons, this procedure is repeated 10000 times with labels (e.g., expected vs. unexpected) being assigned randomly to the data. Using the distribution resulting from those permutations as a null distribution, significance of the found clusters can be established.

### Statistics

Apart from the cluster-based permutations tests (which were carried out in MNE-Python and FieldTrip), all analyses were carried out in JASP (version 0.16.4, 43), with significance established at p < 0.05. If for any of the t-tests the normality assumption was not met, Wilcoxon sign-rank tests were instead carried out. Each frequentist test in JASP was also complemented by a test based on a Bayesian approach (using default uninformed priors): here we compared the evidence for the null model against alternative models using Bayes Factors (BF), allowing us to determine evidence for the presence or absence of an effect (44). For example, a BF of 3 means that the data are three times more likely to be observed under the alternative than the null hypothesis and a BF of 1/3 means that the data are three times more likely to be observed under the null than the alternative hypothesis (conventionally described as providing moderate evidence for the presence or absence of an effect, 44).

## RESULTS

### Preparatory analyses

Initially, we performed several control analyses to assess task performance and establish the presence of all necessary responses. First, all participants included in the final data analysis were able to discriminate repetitions and omissions with sufficient accuracy (overall discrimination accuracy >50% for every participant, mean accuracy for repetitions: 74.0%, mean accuracy for omissions: 87.9%, Fig. 2a). Second, the individually calibrated stimuli were perceived as painful on average, i.e. received a VAS rating > 50 (t_(35)_=2.6, p=0.007, BF_+0_=6.657). Interestingly, pain ratings differed between the experimental conditions as revealed by a two-factorial repeated-measures ANOVA (Fig. 2b): while we observed no main effect of stimulus position (i.e. first stimulus vs. repetition; F_(1,35)_=0.007, p=0.932, BF_incl_ = 0.381, Bayes factor indicates absence of sufficient evidence), we did observe an effect of block type (i.e. high vs. low, with high blocks being rated as more painful than low blocks; F_(1,35)_=29.2, p<0.001, BF_incl_=4.3 × 10^3^), but no significant interaction effect (F_(1,35)_=2.1, p=0.153, BF_incl_=0.492, indicating absence of sufficient evidence). Third, we observed that participants formed the desired expectations (Fig. 2c), i.e. reported to expect significantly less than 50% repetitions in the low blocks (mean=22.0%, W=0, p<0.001, BF_-0_=1.9×10^5^) and more than 50% repetitions in the high blocks (mean=75.1%, t_(35)_=22.8, p<0.001, BF_+0_=5.5×10^19^). Finally, we observed all hypothesised EEG responses (assessing first-stimulus responses only), encompassing the N1 component with a lateralized topography (latency=238ms, amplitude=-1.3µV, Fig. 2d), the N2P2 complex with a central topography (N2: latency=264ms, amplitude=-2.1µV; P2: latency=390ms, amplitude=4.1µV, Fig. 2d), a desynchronization in the alpha and beta band as well as an increase in gamma band oscillations (GBOs; Fig. 2e).

**Figure 2:**
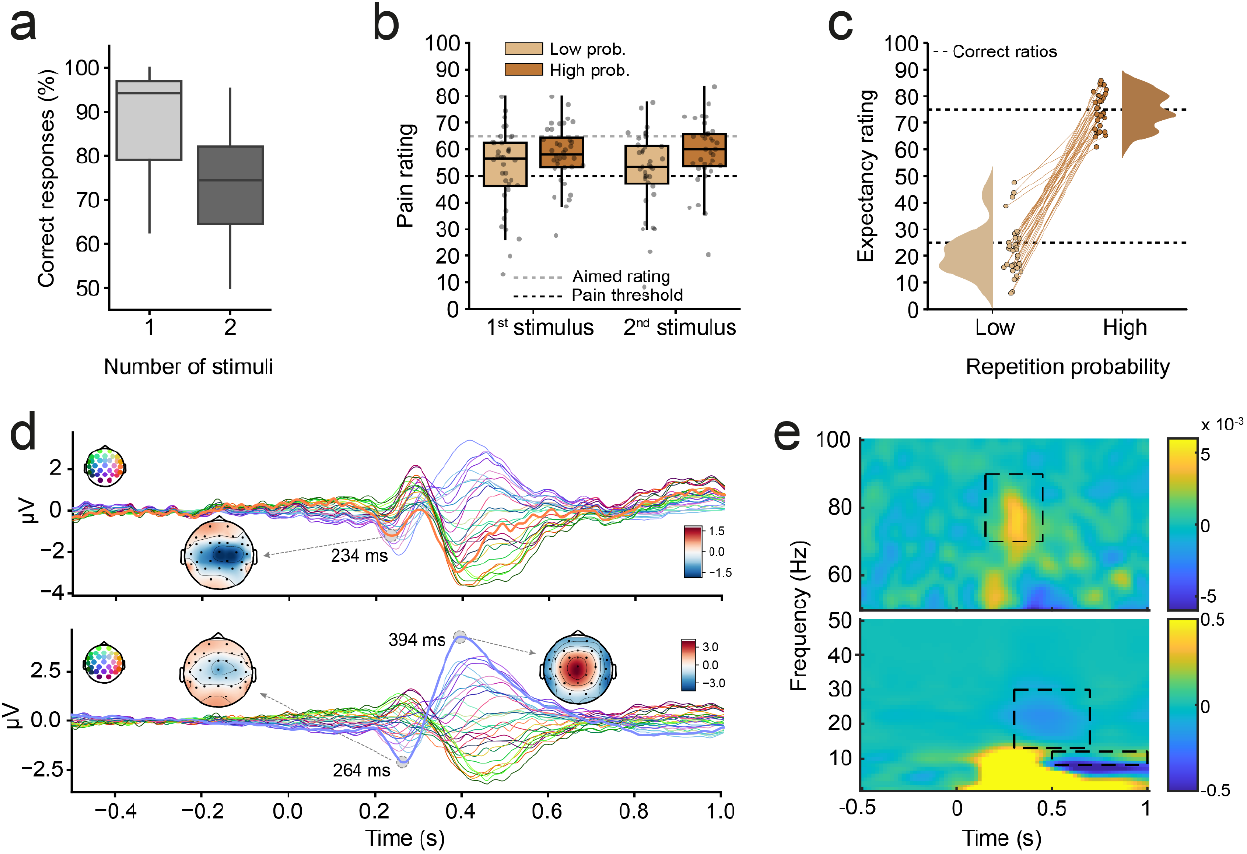
Control analyses. The displayed data encompasses all trials. (a) Percentage of correct responses in the control task across participants, depicted separately for trials with one and two stimuli. (b) Pain ratings, depicted separately for block type (high vs. low block) and stimulus position in trial (first vs. second stimulus), with dotted lines representing the pain threshold and the aimed-for rating. Ratings above 50 describe painful sensations. (c) Expectancy ratings. Distributions and single participant datapoints depicted separately for the block types. Lines connect datapoints per participant and dotted lines represent the correct contingencies (25% for low blocks, 75% for high blocks). (d) LEPs obtained from single electrodes in response to the first stimulus and associated component topographies. Top row: N1 (reference: electrode Fz), bottom row: N2 and P2 (reference: average of all electrodes). Line colors correspond to the electrodes depicted schematically the top left of each panel, with bold lines representing T8-Fz and Cz-avg for N1 and N2P2, respectively. (f) Time-frequency response to first stimuli, absolute power change compared to baseline (−0.75 to −0.25s) at Cz, with dotted regions depicting ROIs (alpha, beta, gamma).

### EEG repetition suppression

We assessed EEG repetition suppression by taking the repetition trials from both block types into account. We observed repetition suppression of the LEPs, with the N1 amplitude decreasing by 21.4% (though this decrease was not significant: W=400, p=0.150, BF_+0_=0.816; Fig. 3a) and the N2P2 peak-to-peak amplitude decreasing by 43.3% (W=649, p<0.001, BF_+0_=4591; Fig. 3b). In the time-frequency domain, only the GBOs showed a significant repetition suppression (W=470, p=0.015, BF_+0_=1.655, Fig. 3c). Alpha (t_(35)_=0.028, p=0.489, BF_+0_=0.183, Fig. 3c) and beta desynchronisations (W=194, p=0.986, BF_+0_=0.061, Fig. 3c) did not show any significant repetition suppression, with Bayes factors providing at least moderate evidence for the absence of repetition suppression.

**Figure 3:**
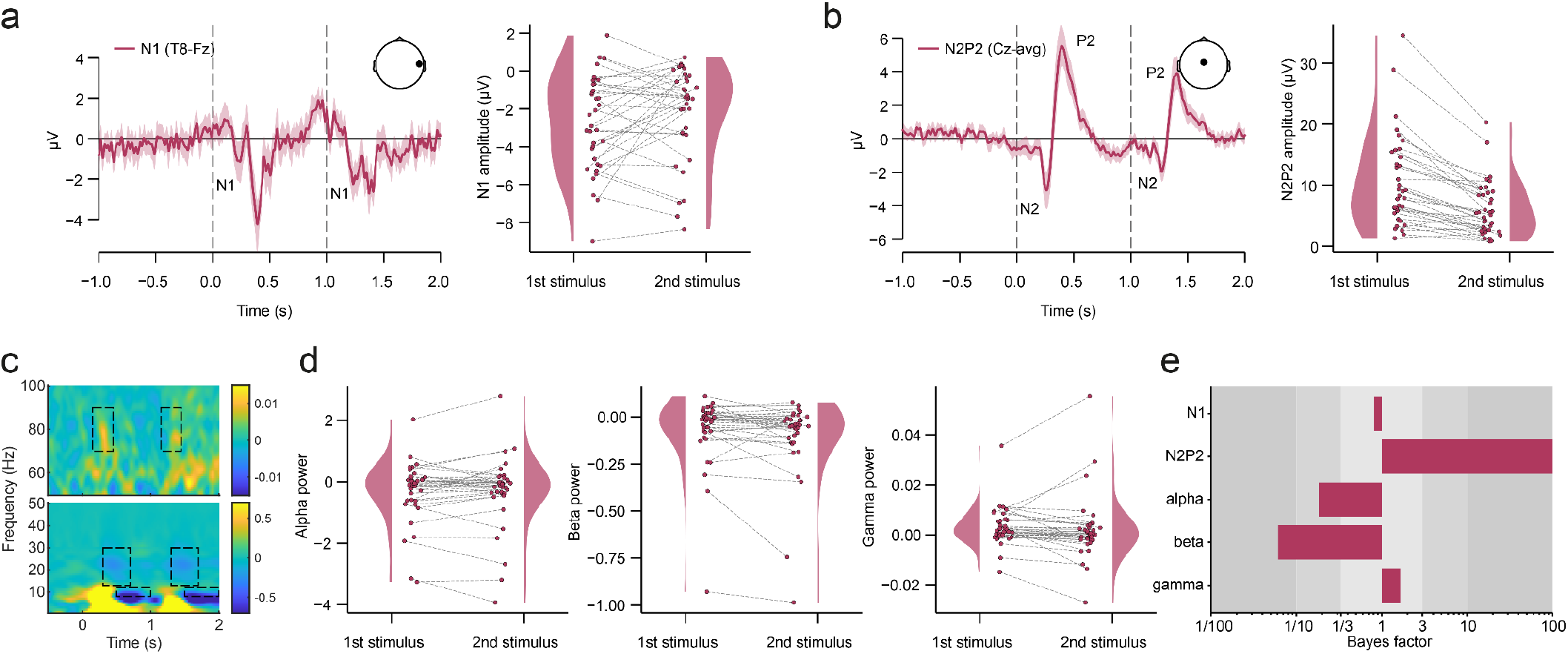
Repetition suppression. The displayed data encompasses only the first half of the experiment with trial counts equalized between conditions. (a) N1 response. The left panel displays the grand average time-course at T8-Fz, with dotted lines depicting stimulus onsets. The right panel shows extracted amplitudes for first and second stimulus per participant. (b) N2P2 response. The left panel displays the grand average time-course at Cz-average, with dotted lines depicting stimulus onsets. The right panel shows extracted amplitudes for first and second stimulus per participant. (c) TFR showing absolute power change compared to the baseline period (− 0.75s to −0.25s), grand average TFR at Cz, with dotted lines depicting ROIs (alpha, beta, gamma). (d) Extracted mean power values from the alpha, beta and gamma ROI, respectively, for first and second stimulus per participant. (e) Bayes factors: dependent-samples Bayesian t-test assessing whether response to second stimulus is smaller than response to first stimulus.

### Expectation effects on repetition suppression in the EEG data

Next, we compared possible amplitude reductions due to repetition between the different blocks (i.e. high vs. low, translating to expected vs. unexpected repetitions, respectively). Neither for the N1 (t_(35)_=0.50, p=0.310, BF_+0_=0.275; Fig. 4a), nor for the N2P2 complex (t_(35)_=-0.558, p=0.710, BF_+0_=0.123; Fig. 4b) did we observe a significant effect of expectations, with Bayes factors providing moderate evidence against an effect of expectations on the EEG repetition suppression magnitude. Similarly, none of the ROIs in the time-frequency domain showed a significant modulation of repetition suppression by expectations, with Bayes factors revealing an absence of sufficient evidence for either the null or the alternative hypothesis (alpha: W=403, p=0.139, BF_+0_=0.447; beta: W=418, p=0.093, BF_+0_=0.563; gamma: W=416, p=0.099, BF_+0_=0.601; Fig. 4c, d), though all Bayes factors pointed towards null effects. The cluster-based permutation tests performed to search for effects outside our predefined response measures (peak amplitudes or time-frequency ROIs) did not reveal any significant clusters in the time domain, and the various small but significant clusters in the time-frequency domain revealed no clear picture, with respect to direction, latency or frequency of effect, especially since the majority was found outside meaningful time windows (i.e. not after the second stimulus).

**Figure 4:**
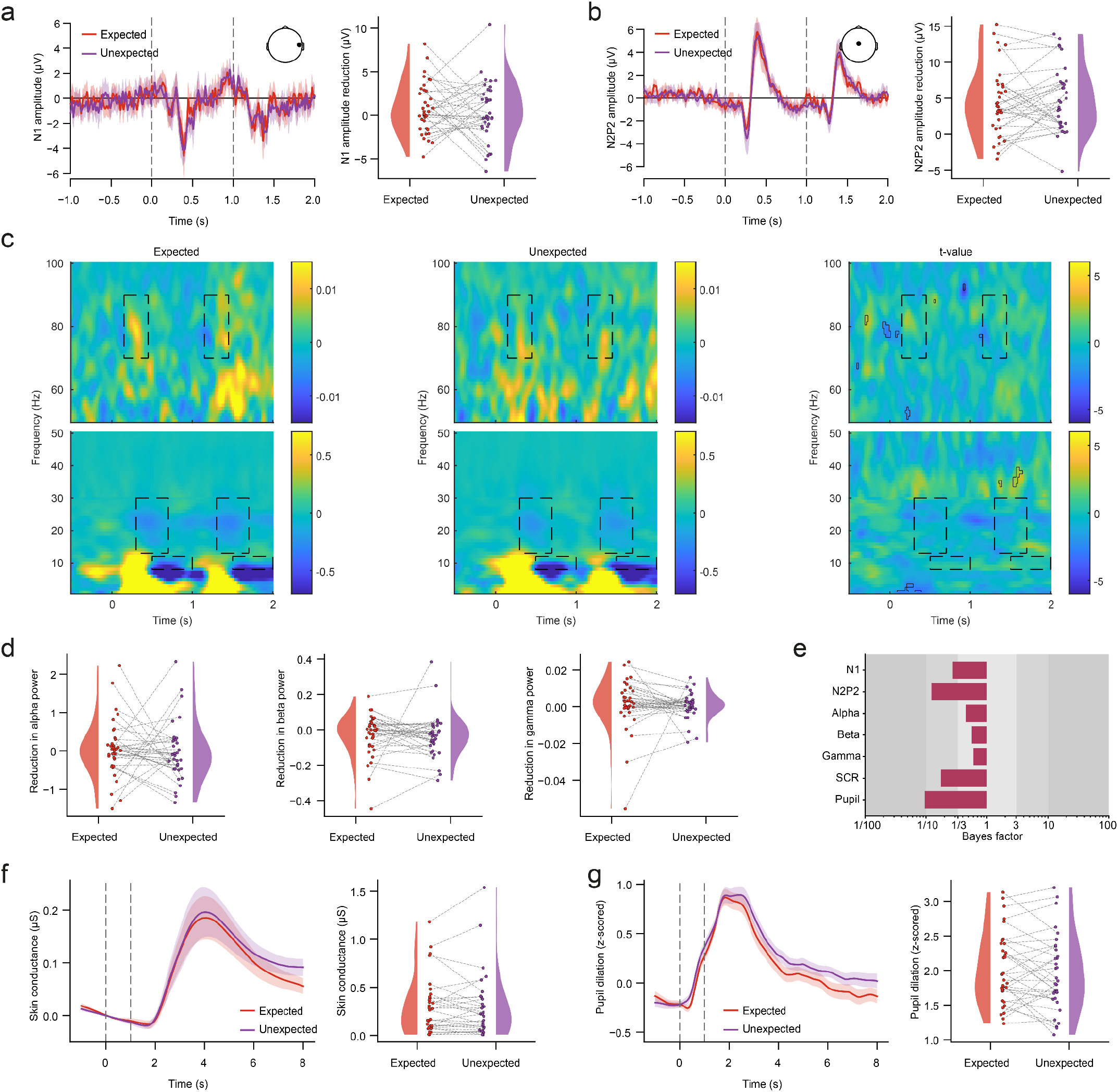
Expectational modulation of repetition suppression. The displayed data encompasses only the first half of the experiment with trial counts equalized between conditions. (a) N1 response. The left panel displays the grand average time-course of expected and unexpected repetitions at T8-Fz, with dotted lines depicting stimulus onsets. The right panel shows extracted amplitudes for expected and unexpected repetitions per participant. (b) N2P2 response. The left panel displays the grand average time-course of expected and unexpected repetitions at Cz-average, dotted lines depict stimulus onsets. Right panel shows extracted amplitudes for expected and unexpected repetitions per participant. (c) TFR showing absolute power change compared to the baseline period (−0.75s to −0.25s) at electrode Cz. The left panel displays grand average TFR of unexpected repetition trials, the middle panel displays grand average TFR of expected repetition trials, and the right panel displays a t-value map of differences. Dotted lines depict ROIs (alpha, beta, gamma) and solid black lines depict significant clusters of the cluster-based permutation test. (d) Extracted mean power values from the TFR ROIs for expected and unexpected repetitions. The left, middle and right panels show alpha, beta and gamma power, respectively. (e) Bayes factors of dependent-samples Bayesian t-test assessing whether repetition suppression is stronger for expected than unexpected repetitions (for N1, N2P2, alpha, beta and gamma) or whether the combined response to expected repetition trials is smaller than to unexpected repetitions (for SCR and pupil dilation). (f) SCR. The left panel displays the grand average time-course of skin conductance responses for expected and unexpected repetitions, with dotted lines depicting stimulus onsets. The right panel shows extracted amplitudes for expected and unexpected repetitions per participant. (g) Pupil dilation. The left panel displays grand average timecourse of pupil dilation for expected and unexpected repetitions, with dotted lines depicting stimulus onsets. The right panel shows extracted amplitudes for expected and unexpected repetitions per participant.

### Expectation effects on repetitions in peripheral physiological data

For both skin conductance and pupil dilation responses, we compared the peak amplitudes in response to a repeated stimulus between high and low blocks, translating to expected and unexpected repetitions. The tests revealed moderate to strong evidence against an expectation effect on the amplitude in repetition trials in both modalities (skin conductance: t_(30)_=-0.116, p=0.546, BF_+0_=0.176; Fig. 4f; pupil dilation: t_(33)_=-1.084, p=0.857, BF_+0_=0.095; Fig. 4g). Additionally, we tested the stimulus time courses within the whole interval after the laser stimuli for any differences between the unexpected and expected condition using a cluster-based permutation test. Although both measures showed slightly higher traces in the unexpected condition, no significant clusters were observed.

## DISCUSSION

In this study, we assessed the mechanisms that underlie a fundamental feature of sensory processing – repetition suppression – in the nociceptive domain. Specifically, we aimed to distinguish between bottom-up and top-down explanations for repetition suppression to nociceptive stimuli by assessing the influence expectations exert on repetition suppression metrics in healthy volunteers. We modified an established repetition suppression paradigm from the auditory domain and employed painful CO_2_ laser stimuli for selective nociceptive stimulation. We recorded subjective ratings, autonomic nervous system responses as well as EEG time-domain and time-frequency domain responses and analysed the data with both frequentist and Bayesian approaches, aiming to provide a comprehensive picture of mechanisms of repetition suppression in nociception.

### Responses to nociceptive laser stimulation across processing levels

We first ensured – via extensive calibration procedures and thereby informed subsequent participant exclusion – that the here-employed CO_2_-laser stimuli were perceived as painful. This was deemed necessary considering the well-established correlation of gamma-band oscillations with subjectively perceived pain intensity (25–27). The applied nociceptive stimulation also led to robust responses in the autonomic nervous system recordings we performed, i.e. we were able to detect clear laser-evoked skin conductance as well as pupil dilation responses. With regards to EEG responses, we observed robust responses in the time-domain (i.e. N1 and N2P2 potentials) as well as in the time-frequency domain (i.e. stimulus-induced alpha and beta desynchronization as well as increased gamma band oscillations, GBOs). By using i) a thorough denoising procedure to remove muscle artifacts (35) and ii) a subtractive rather than a divisive baseline correction to prevent an artificial increase of synchronisations (36), we aimed to eliminate artefactual contributions to the observed GBOs.

In all response measures, we observed a substantial habituation over the course of the experiment and hence decided to base all analyses on the first half of the experiment only. Such habituation is not uncommon especially for the N2P2 component (as well as the N1 component, 45). To our knowledge though, the long-term habituation of oscillatory components to nociceptive stimulation is a novel finding, as studies investigating habituation at the timescale of this experiment (approximately one hour) do not exist. Such habituation might be addressed in future studies by adding catch-trials to enhance participants’ vigilance over the course of the entire experiment.

### Repetition suppression

With regards to repetition suppression (i.e. a reduced response to the second stimulus of a pair), we observed a substantial effect for the N2P2 complex (>40% reduction), which is well in line with studies using similar repeated laser stimuli in EEG recordings (23,24,46–49) and ECoG recordings (50) as well as repetition suppression in other sensory domains (7,9,51). Repetition suppression of the N1 component was also observed (>20%), but failed to reach significance, which fits with a broader picture of habituation in nociception, where habituation of the N1 component has been observed less frequently (45).

Importantly, we found a significant repetition suppression of vertex-recorded GBOs, which is in line with the idea of GBOs coding prediction errors (21) as well as ECoG recordings from the insula showing habituation of GBOs to triplet stimulation (50). However, these results are in contrast to an EEG study using a similar paradigm (28) where no GBO suppression was observed. GBOs in the latter study however spanned a broader range of frequencies and exhibited a more lateral topography. Taken together, it is likely that different processes underlie these divergent GBO results (see also discussion in Liberati et al., 50) and full elucidation of the different components of laser-induced GBOs awaits further studies, with recent progress being made in animal models (52).

In contrast to the repetition suppression for N1, N2P2 and GBOs, we did not observe repetition suppression of low frequency desynchronisations in the alpha and beta range. These findings complement and extend previous evidence (46) by providing evidence for the absence of such an effect via Bayesian analyses. While the functional significance of these cortical dissociations is currently unclear, they clearly indicate multiple – and possibly opposing (see also 45) – processes, and thus not a simple and homogeneous feed-forward relay from lower levels. It is also highly unlikely that the observed repetition suppression effects result from adaptation effects at peripheral levels, since the laser beam was always shifted between the two stimuli of a pair.

Interestingly, the observed repetition suppression in LEPs and GBOs was not mirrored on the perceptual level, i.e. Bayesian analyses provided evidence that participants rated first and second stimuli as similarly intense. This dissociation between neural and subjective effects could have several reasons. First, LEPs are not necessarily indicators of nociceptive processing per se, but rather reflect saliency-related processes and are thus not suitable as a readout for subjective pain perception (46,53). While GBOs have displayed a closer link to the perceptual level (25,26,28,54,but see 50), the dissociation between GBOs and pain ratings could arise from the different recording timescales: all EEG metrics were based on trial-wise recordings, whereas pain ratings could only be obtained retrospectively after each block and memory effects could thus play a substantial role here (45). Further, since performance in the choice discrimination task for dissociating one vs two stimuli was not at ceiling level there might have been some difficulty in retrospectively rating two distinct percepts precisely.

### Expectation effects on repetition suppression

Most importantly, we aimed to assess whether the observed repetition suppression was driven by the fact that the second stimulus was more predictable than the first one. We therefore varied the expectational context in which repetitions were presented across blocks. We observed that despite participants forming the correct expectations as indicated by their pre-block expectation ratings, none of the outcome measures, neither time-domain and time-frequency domain EEG nor autonomic nervous system measures showed any significant modulation of repetition suppression by expectations; this was reinforced by the results of Bayesian analyses providing rather consistent evidence for an absence of such effects. This stands in direct contrast to our hypothesis and also to results from a study in the auditory domain upon which we based our paradigm (5) and where unexpected repetitions showed much lower suppression, particularly in GBOs.

It does however tie in with, and add to, previous studies in the nociceptive domain, where expectations regarding *location* or *modality* were manipulated and no EEG repetition suppression was observed (23,24,but see 22, where in one of four conditions the temporal predictability had a significant effect). We extend these previous findings by showing that repetition suppression is also not influenced by the most fundamental expectation manipulation, namely that of *repetition probability*. Additionally, in contrast to all previous repetition suppression studies in the nociceptive domain we also included GBOs in our outcome metrics – as these had been linked to prediction errors in other sensory domains (29,for an example in heat pain see 55) – and also did not observe a modulation by expectations, further reinforcing the lack of expectation effects in EEG metrics. It is noteworthy however that there has been a recent report of dissociation between local oscillations (which we investigate here) and interregional connectivity with respect to sensory and expectation effects in the nociceptive domain (56).

Regarding the reasons for the observed lack of expectational modulation, at least three aspects deserve to be mentioned. *First and foremost*, predictive coding models suggest that to guide our interpretation of the world optimally, the precision of both prior knowledge and sensory input needs to be considered (57–59). The more precise our predictions about the external world are and the more ambiguous the sensory input is, the stronger the weighing of top-down factors, such as expectation, will be. Considering however that we created highly precise sensory input (by employing strong and short laser stimulation), this might lead to a higher weight associated with the sensory input and thus bias the cortical responses assessed here towards bottom-up input (37,60). Therefore, adaptation processes – in contrast to expectation processes – might be the prevalent mechanism for the repetition suppression observed here. Supporting this idea are data from the auditory domain (61): when trying to quantify the proportions of expectation-independent adaptation processes and prediction error reduction in repetition suppression, it has been shown that this proportion is more shifted towards prediction errors if stimuli are less intense, as predictions are then more relevant to interpret the sensory input meaningfully. *Second*, although participants rated their pre-block expectations in the hypothesized way, such ratings can obviously be influenced by desirability (62) and might thus not be a behavioural read-out of predictions. *Finally*, a fundamental difference between our study and data from experiments with visual or auditory stimuli (e.g., 5) is the different temporal nature of input in these domains: the visual and auditory systems in humans typically receive a rather constant stream of input and thus a sparse encoding and suppression of unchanging elements can be argued to be necessary for novelty detection and adaptive behaviour. The human nociceptive system on the other hand usually receives phasic input and thus the need for suppression of unchanging background activity is far less common (although arguments have also been presented for nociception to be a more continuously operating process than traditionally assumed when considering the subjective experience of pain (63). In this regard it is also important to note that descending pain modulation systems – which have been argued to carry predictions (60) – show a selectivity in modulation and often leave A-fibre mediated sensory-discriminative processing, underlying the short-latency EEG responses reported here, intact (64).

### Limitations

Several limitations of this study deserve to be mentioned. First, while the stimulus discrimination task showed participants being able to distinguish single-from double-stimulation, their performance was not at ceiling level. This might be improved with shorter stimulus duration as provided for example by Nd:YAP lasers, which could also have the added benefit of a higher signal-to-noise ratio in the EEG responses due to a more synchronous afferent volley. However, this might not necessarily lead to more prominent expectation effects due to the increased precision of the sensory input. Second, due to the substantial habituation we observed and the ensuing focus on the first half of the study, the number of trials per condition was relatively small and statistical power thus rather low. Finally, we only investigated one electrophysiological processing level, i.e. cortical LEPs and oscillations, which limits the inferences we can draw, especially when taking into account that the relative contributions of adaptation and prediction errors to repetition suppression have been shown to differ along the auditory pathway (61). Therefore, to answer open questions about the relevant hierarchical levels for possible expectational modulations, future studies should evaluate responses on as many different processing levels as possible, e.g. including cortical connectivity measures (56) as well as spinal cord responses (65,66).

## Conclusion

In summary, we were able to demonstrate EEG repetition suppression effects in response to nociceptive laser stimuli and observed that repetition suppression was not modulated by expectations regarding repetition probability, favouring bottom-up over top-down explanations for repetition suppression. By employing different outcome measures, thorough data cleaning as well as frequentist and Bayesian statistics, we were able to draw a comprehensive picture regarding the absence of such expectation effects. Whether this pattern of results is dependent on the employed paradigm and the highly precise laser stimulation or indicates a specialized role for nociception - in contrast to other sensory domains – remains a topic for future research.

## Author contributions (according to CRediT taxonomy, https://credit.niso.org)

Conceptualization: FE, UH, LGP, MP

Data curation: LGP

Formal analysis: UH, MMN, LGP

Funding acquisition: FE

Investigation: LGP

Methodology: FE, UH, BN, LGP

Project administration: FE, UH, LGP

Resources: FE

Software: UH, LGP

Supervision: FE, UH

Visualization: FE, UH, LGP

Writing – original draft: FE, UH, LGP

Writing – review & editing: FE, UH, MMN, BN, MP, LGP

## ACKNOWLEDGEMENTS

We would like to thank all volunteers who participated in this study. Additionally, we want to thank our research assistants and interns who assisted in data acquisition and piloting: Pia Baisch, Lukas Feuring, Melanie Freund, Magdalena Gruner, Leonie Heinicke, Duygu Işik, Paula Kosel, Mareike Pauly, Kathrin Warmbier. This manuscript will be part of a doctoral thesis.

