## Supplementary Material for "Expectation effects on repetition suppression in nociception"

### Supplementary results

The supplementary results contain two further sets of analyses. First, although not a primary aim of the study, we investigated omission responses in the EEG and peripheral physiological data. Second, we reran all previous analyses (including the here-presented omission responses) with inclusion of all eight blocks of data – please note that in the main manuscript we focussed only on the first four blocks due to substantial habituation across the experiment.

*Omission responses in EEG data.* We compared responses to expected and unexpected omissions at the timepoints where one would expect stimulus-related responses (i.e. N1, N2P2, alpha, beta and gamma ROIs). We did not observe any significant differences between the conditions, with Bayes factors consistently providing moderate evidence against an expectation effect (N1: $t_{(35)}=-0.554$, $p=0.583$, $BF_{10}=0.207$; N2P2: $t_{(35)}=-0.949$, $p=0.349$, $BF_{10}=0.272$; alpha: $W=360$, $p=0.681$ $BF_{10}=0.188$; beta: $W=388$, $p=0.396$, $BF_{10}=0.281$; gamma: $W=376$, $p=0.509$, $BF_{10}=0.225$; Fig. S1a-e). The additional cluster-based permutation tests did not reveal any significant clusters when comparing expected and unexpected omissions in the time domain, while clusters formed in the time-frequency domain did not reveal a clear picture since they were not restricted to the time after the omission.

*Expectation effects on omissions in peripheral physiological data.* For both skin conductance and pupil dilation responses, we compared the peak amplitudes in response to a single stimulus between high and low blocks, translating to unexpected and expected omissions. The test for skin conductance data revealed evidence against an expectation effect on the amplitude in omission trials while the test for pupil dilation data remained inconclusive (skin conductance: $t_{(30)}=-0.914$, $p=0.816$, $BF_{+0}=0.108$; pupil dilation: $t_{(33)}=1.059$, $p=0.149$, $BF_{+0}=0.519$; Fig. S1f, g). Additionally, we tested the single stimulus time courses within the whole interval after the omission of the laser stimulus for any differences between the unexpected and expected condition using a cluster-based permutation test. For skin conductance time courses no significant cluster was observed, whereas for pupil dilation one significant cluster was observed after the main response ($p=0.0293$, latency: 5.2-7.6s, Fig. S1f) with lower pupil dilation in trials with expected omissions.

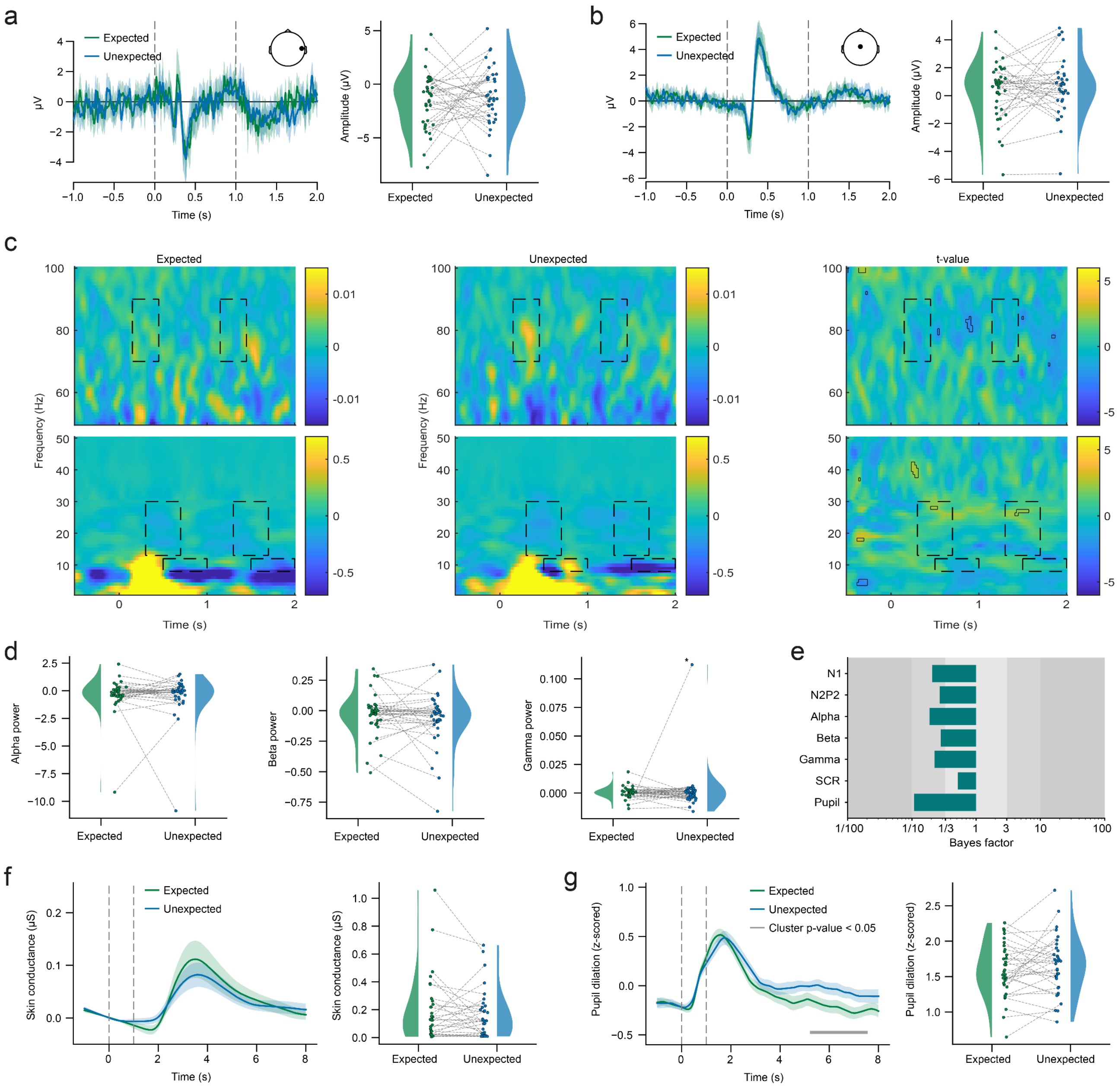


**Figure S1:** Expectation effects on omission responses. The displayed data encompasses only the first half of the experiment with trial counts equalized between conditions. (a) N1 response. The left panel displays the grand average time-course of expected and unexpected omissions at T8-Fz, with dotted lines depicting stimulus onsets. The right panel shows extracted amplitudes for expected and unexpected omissions per participant. (b) N2P2 response. The left panel displays the grand average time-course of expected and unexpected omissions at Cz-average, with dotted lines depicting stimulus onsets. The right panel shows extracted amplitudes for expected and unexpected omissions per participant. (c) TFR showing absolute power change compared to the baseline period (-0.75s to -0.25s) at electrode Cz. The left panel displays grand average TFR of expected omission trials, the middle panel displays grand average TFR of unexpected omission trials, and the right panel displays a t-value map of differences. Dotted lines depict ROIs (alpha, beta, gamma) and solid black lines depict significant clusters of the cluster-based permutation test. One outlier excluded in display (marked in d). (d) Extracted mean power values from the TFR ROIs for expected and unexpected omissions. The left, middle and right panels show alpha, beta and gamma power, respectively. (e) Bayes factors of dependent-samples Bayesian t-test assessing whether responses to expected and unexpected omissions differ (for N1, N2P2, alpha, beta and gamma) or whether the combined response to expected omission trials is smaller than to unexpected omissions (for SCR and pupil dilation). (f) SCR. The left panel displays the grand average time-course of skin conductance responses for expected and unexpected omissions, with dotted lines depicting stimulus onsets. The right panel shows extracted amplitudes for expected and unexpected omissions per participant. (g) Pupil dilation. The left panel displays the grand average time-course of pupil dilation for expected and unexpected omissions, with dotted lines depicting stimulus onsets; the grey bar below the time-courses marks a significant cluster of the cluster-based permutation test. The right panel shows extracted amplitudes for expected and unexpected omissions per participant.

*Entire dataset: EEG repetition suppression*. We evaluated EEG repetition suppression by considering repetition trials from both types of blocks. Our analysis revealed repetition suppression of the LEPs, with a reduction of the N1 amplitude by 11.7% (though this reduction was not significant: $t_{(35)}=0.775$, $p=0.222$, $BF_{+0}=0.365$; Fig. S2a) and of the N2P2 peak-to-peak amplitude by 38.1% ($t_{(35)}=6.315$, $p<0.001$, $BF_{+0}=10.7e4$; Fig. S2b). In the time-frequency domain, no significant repetition suppression was observed (alpha: W=182, p=0.992, $BF_{+0}$=0.062; beta: W=110, p=1.000, $BF_{+0}$=0.075; gamma: W=421, p=0.086, $BF_{+0}$=0.608, Fig. S2c), with most Bayes factors indicating evidence for the absence of repetition suppression.

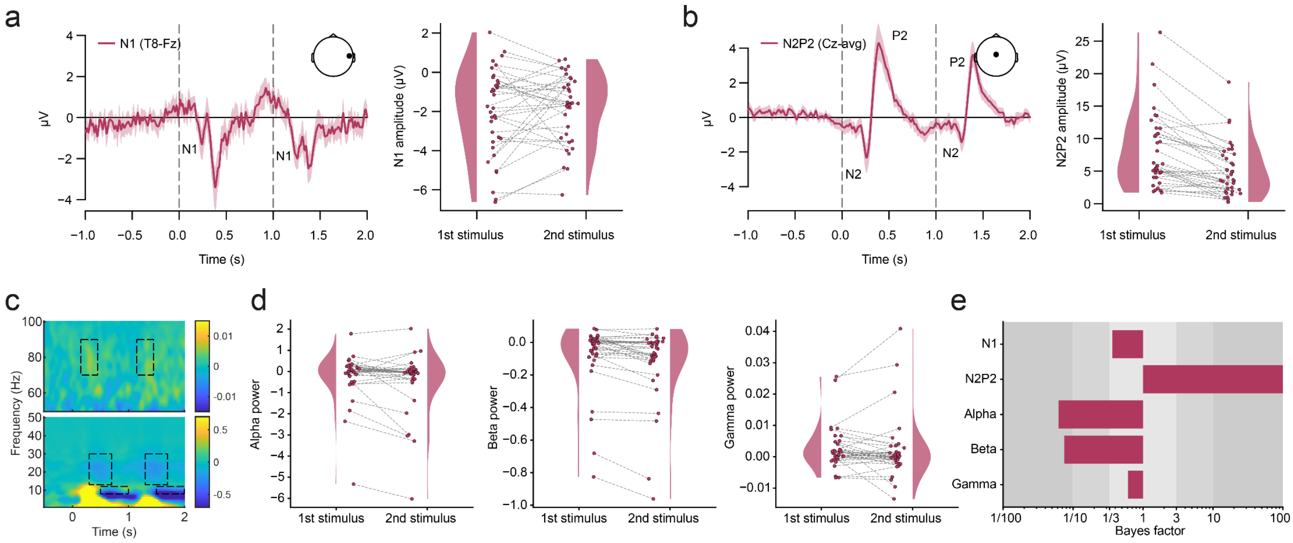


**Figure S2:** Repetition Suppression. The displayed data encompasses data from the whole experiment with trial counts equalized between conditions. (a) N1 response. The left panel displays the grand average time-course at T8-Fz, with dotted lines depicting stimulus onsets. The right panel shows extracted amplitudes for first and second stimulus per participant. (b) N2P2 response. The left panel displays the grand average time-course at Cz-average, with dotted lines depicting stimulus onsets. The right panel shows extracted amplitudes for first and second stimulus per participant. (c) TFR showing absolute power change compared to the baseline period (-0.75s to -0.25s), grand average TFR at Cz, with dotted lines depicting ROIs (alpha, beta, gamma). (d) Extracted mean power values from the alpha, beta and gamma ROI, respectively, for first and second stimulus per participant. (e) Bayes factors: dependent-samples Bayesian t-test assessing whether response to second stimulus is smaller than response to first stimulus.

*Entire dataset: expectation effects on repetition suppression in the EEG data.* Subsequently, we tested for an expectational modulations of possible amplitude reductions due to repetition by contrasting responses from high and low blocks (corresponding to expected vs. unexpected repetitions, respectively). Our analysis did not reveal a significant effect of expectations on the N1 component ($t_{(35)}$=0.070, p=0.472, $BF_{+0}$=0.189; Fig. S3a) or the N2P2 complex ($t_{(35)}$=0.143, p=0.444, $BF_{+0}$=0.200; Fig. S3b), with Bayes factors indicating moderate evidence against the influence of expectations on the magnitude of EEG repetition suppression. Surprisingly, in the time-frequency domain we observed a significant effect of expectations on repetition suppression of alpha and beta desynchronizations, however Bayes factors remain inconclusive (alpha: W=467, p=0.017, $BF_{+0}$=1.898; beta: $t_{(35)}$=1.972, p=0.028, $BF_{+0}$=1.957; Fig. S3c, d). When comparing the mean power reduction values, it however became obvious that this significant effect results from a stronger desynchronization for the repetition compared to the first stimulus, i.e. a repetition enhancement. This enhancement is significantly stronger for unexpected compared to expected repetitions. The gamma band ROI did not show a significant effect of expectations on repetition suppression, with an inconclusive Bayes factor (W=402, p=0.143, $BF_{+0}$=0.373; Fig. S3c, d). Additional cluster-based permutation tests failed to identify significant clusters within the time domain. Furthermore, the small yet statistically significant clusters observed in the time-frequency domain did not present a coherent pattern regarding the direction, latency, or frequency of the effects with most clusters being detected outside meaningful time windows (before the second stimulus).

*Entire dataset: expectation effects on repetitions in peripheral physiological data.* To investigate expectation effects on both skin conductance and pupil dilation responses, peak amplitudes for repeated stimuli were compared between high and low blocks (corresponding to expected and unexpected repetitions). The results yielded moderate to strong evidence against an expectation effect on repetition trial amplitudes in both modalities (skin conductance: $t_{(30)}$=-0.962, p=0.828, $BF_{+0}$=0.106; Fig. S3f; pupil dilation: $t_{(33)}$=-2.044, p=0.976, $BF_{+0}$=0.066; Fig. S3g). Furthermore, we employed a cluster-based permutation test to evaluate the entire interval following the laser stimuli for any differences between the unexpected and expected conditions. No significant clusters were observed.

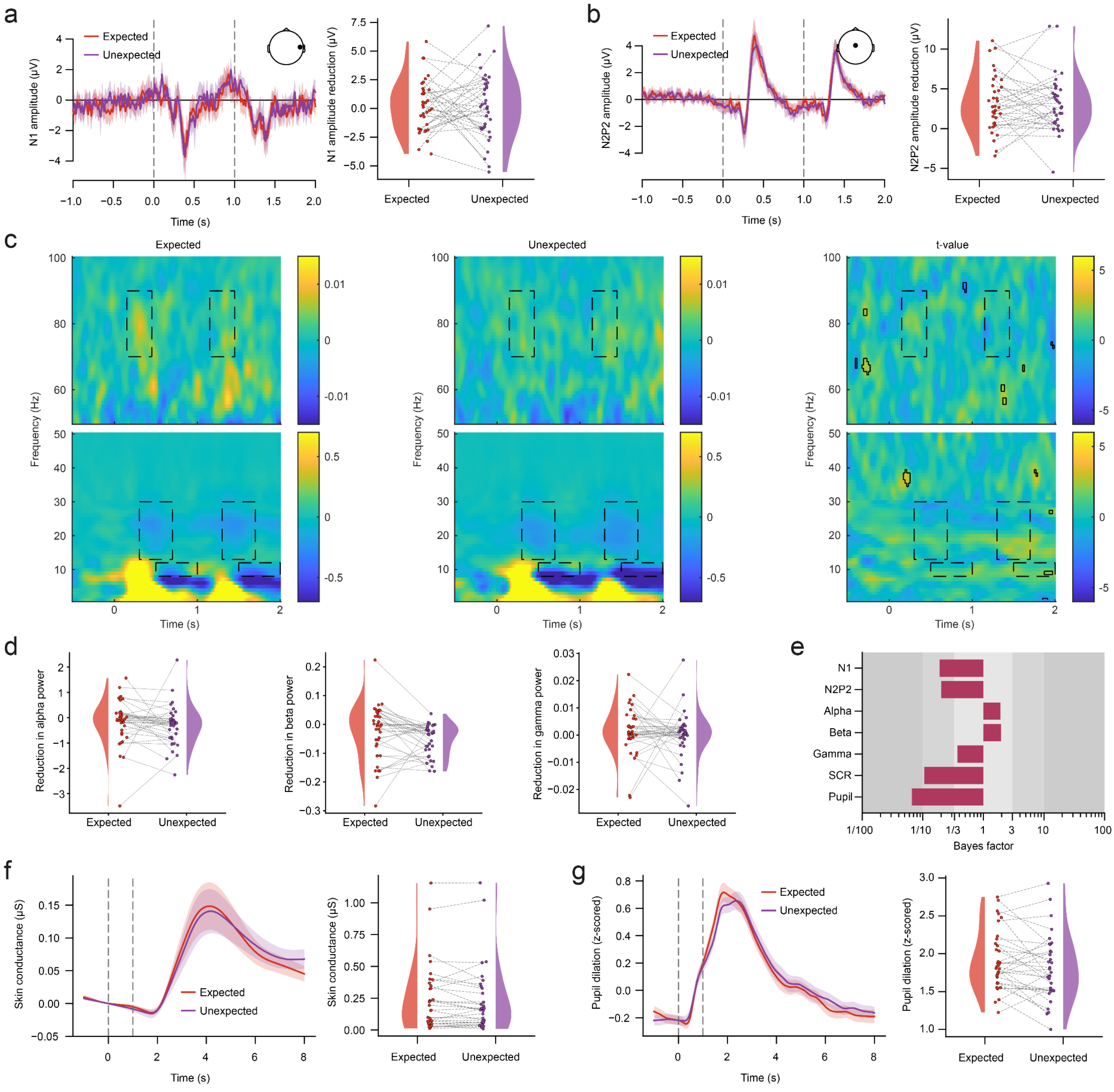


**Figure S3**: Expectation effects on repetition suppression. The displayed data encompasses data from the whole experiment with trial counts equalized between conditions. (a) N1 response. The left panel displays the grand average time-course of expected and unexpected repetitions at T8-Fz, with dotted lines depicting stimulus onsets. The right panel shows extracted amplitudes for expected and unexpected repetitions per participant. (b) N2P2 response. The left panel displays the grand average time-course of expected and unexpected repetitions at Cz-average, with dotted lines depicting stimulus onsets. The right panel shows extracted amplitudes for expected and unexpected repetitions per participant. (c) TFR showing absolute power change compared to the baseline period (-0.75s to -0.25s) at electrode Cz. The left panel displays grand average TFR of unexpected repetition trials, the middle panel displays grand average TFR of expected repetition trials, and the right panel displays a t-value map of differences. Dotted lines depict ROIs (alpha, beta, gamma) and solid black lines depict significant clusters of the cluster-based permutation test. (d) Extracted mean power values from the TFR ROIs for expected and unexpected repetitions. The left, middle and right panels show alpha, beta and gamma power, respectively. (e) Bayes factors of dependent-samples Bayesian t-test assessing whether repetition suppression is stronger for expected than unexpected repetitions (for N1, N2P2, alpha, beta and gamma) or whether the combined response to expected repetition trials is smaller than to unexpected repetitions (for SCR and pupil dilation). (f) SCR. The left panel displays the grand average time-course of skin conductance responses for expected and unexpected repetitions, with dotted lines depicting stimulus onsets. The right panel shows extracted amplitudes for expected and unexpected repetitions per participant. (g) Pupil dilation. The left panel displays the grand average time-course of pupil dilation for expected and unexpected repetitions, with dotted lines depicting stimulus onsets. The right panel shows extracted amplitudes for expected and unexpected repetitions per participant.

*Entire dataset: omission responses in the EEG data.* Additionally, we evaluated the difference between responses to expected and unexpected omissions by extracting the amplitudes of mean power values at the timepoints associated with stimulus-related responses (i.e. N1, N2P2, alpha, beta and gamma ROIs). The results indicated no significant differences between the conditions, with Bayes factors predominantly offering moderate evidence against the presence of an expectation effect (N1: $t_{(35)}=1.773$, $p=0.085$, $BF_{10}=0.735$; N2P2: $t_{(35)}=0.328$, $p=0.744$, $BF_{10}=0.188$; alpha: $W=327$, $p=0.932$, $BF_{10}=0.185$; beta: $W=304$, $p=0.658$, $BF_{10}=0.191$; gamma: $W=329$, $p=0.957$, $BF_{10}=0.195$; Fig. S4a-e). Moreover, cluster-based permutation tests did not identify any significantly different clusters between expected and unexpected omissions in the time domain, and clusters identified in the time-frequency domain lacked clarity as they were not confined to the post-omission period.

*Entire dataset: expectation effects on omissions in peripheral physiological data.* Finally, we assessed the peak amplitudes elicited by a single stimulus for high and low blocks, corresponding to unexpected and expected omissions, for both skin conductance and pupil dilation responses. We found moderate to strong evidence against an expectation effect on the amplitude in omission trials (skin conductance: $t_{(30)}=-1.997$, $p=0.973$, $BF_{+0}=0.070$; pupil dilation: $t_{(33)}=0.372$, $p=0.356$, $BF_{+0}=0.251$). Moreover, we investigated the time courses of the single stimuli within the entire interval following the laser stimulus omission for any differences between the unexpected and expected conditions using a cluster-based permutation test, which did not identify any significant clusters.

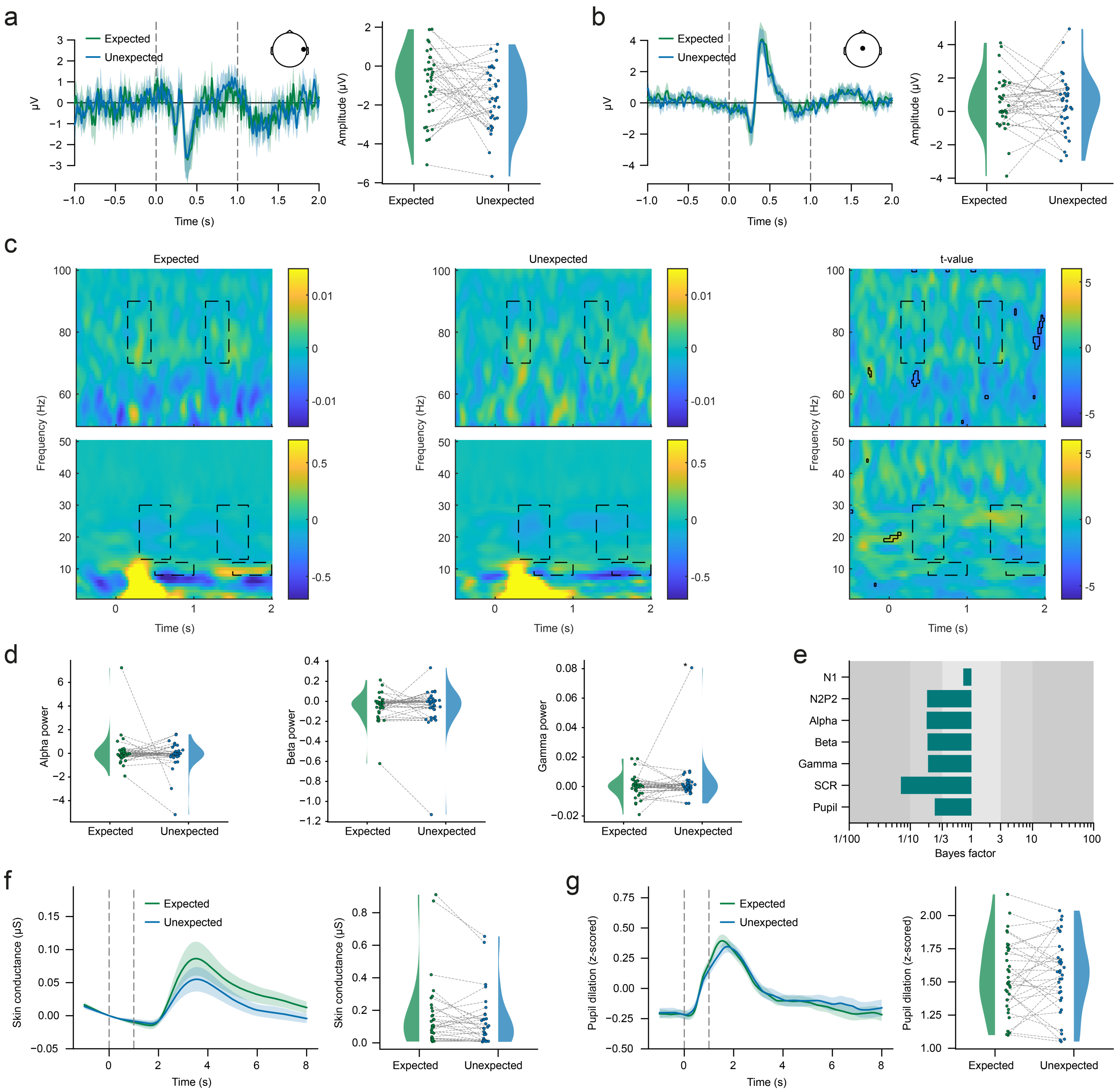


**Figure S4:** Expectation effects on omission responses. The displayed data encompasses data from the whole experiment with trial counts equalized between conditions. (a) N1 response. The left panel displays the grand average time-course of expected and unexpected omissions at T8-Fz, with dotted lines depicting stimulus onsets. The right panel shows extracted amplitudes for expected and unexpected omissions per participant. (b) N2P2 response. The left panel displays the grand average time-course of expected and unexpected omissions at Cz-average, with dotted lines depicting stimulus onsets. The right panel shows extracted amplitudes for expected and unexpected omissions per participant. (c) TFR showing absolute power change compared to the baseline period (-0.75s to -0.25s) at electrode Cz. The left panel displays grand average TFR of expected omission trials, the middle panel displays grand average TFR of unexpected omission trials, and the right panel displays a t-value map of differences. Dotted lines depict ROIs (alpha, beta, gamma) and solid black lines depict significant clusters of the cluster-based permutation test. One outlier excluded in display (marked in d). (d) Extracted mean power values from the TFR ROIs for expected and unexpected omissions. The left, middle and right panels show alpha, beta and gamma power, respectively. (e) Bayes factors of dependent-samples Bayesian t-test assessing whether responses to expected and unexpected omissions differ (for N1, N2P2, alpha, beta and gamma) or whether the combined response to expected omission trials is smaller than to unexpected omissions (for SCR and pupil dilation). (f) SCR. The left panel displays the grand average time-course of skin conductance responses for expected and unexpected omissions, with dotted lines depicting stimulus onsets. The right panel shows extracted amplitudes for expected and unexpected omissions per participant. (g) Pupil dilation. The left panel displays the grand average time-course of pupil dilation for expected and unexpected omissions, with dotted lines depicting stimulus onsets. The right panel shows extracted amplitudes for expected and unexpected omissions per participant.

### Supplementary discussion

*Assessing omission responses.* In addition to investigating repetition suppression, the chosen paradigm also allowed for an investigation of those trials in which the stimulus was expected, but not repeated, i.e. omission trials. Contrary to predictive coding assumptions, we did not observe differences between expected and unexpected omissions in any of the above-mentioned outcome measures, except for a very late difference in the pupil dilation data (more than five seconds after the first stimulus); Bayesian analyses were consistently in favour of no differences. This contrasts strongly with a study from the auditory domain upon which we based our paradigm (1), where significant GBOs were observed for unexpected omissions, and also with studies on omission responses from different sensory domains (auditory: 2, for a review see 3, visual: 4,5, somatosensory: 6–8). It is important to note however, that to our knowledge electrophysiological omission responses have not been investigated in the nociceptive domain so far (though there is some evidence from fMRI studies, e.g., 9,10). Further, since the main goal of this study was to investigate repetition suppression, our paradigm was not optimised for the detection of omission responses, thus the maximum of 12 unexpected omissions was likely too low to observe robust omission responses.

*Differences between the results using the first half of the data versus the full dataset.* While the overall pattern of results was very similar when comparing the results obtained from the first half of the experiment (main manuscript) to those obtained from the full dataset (supplement), there were two main differences, which will be discussed in the following section. First, we did not observe a significant repetition suppression of GBOs in the full dataset, although this was present in the first half of the data. One reason for this lack of repetition suppression could be the strong habituation across blocks (refer to the section "Methods – Preparatory analysis"), suggesting that a further decrease in GBOs through repetition suppression could become undetectable due to the reduced SNR. Such a lack of repetition suppression with very high overall stimulus numbers was also observed by Zhang et al. (11). Please note that a comparison with repetition suppression of GBOs in ECoG recordings of the insula is not possible, since no information of total trial counts was available (12). Second, in the full dataset we found a significant expectational modulation of repetition suppression in the alpha and beta band ROIs, which was not present in the first half of the dataset. Canonical responses in both ROIs are desynchronizations (13) and thus repetition suppression becomes evident in higher power values as a response to the repeated stimulus, i.e. less desynchronization. To account for this, we calculated repetition suppression as the difference between the response to the second stimulus and the response to the first stimulus, so that positive values describe a reduction of post-stimulus desynchronization. However, a more thorough examination of the values in the full dataset revealed a contrasting pattern: desynchronizations are stronger after a repetition. This repetition enhancement is not captured by our quantification of repetition suppression, as we conducted one-sided tests exclusively due to our strong a-priori hypothesis. The comparison of this repetition enhancement between the two expectancy conditions revealed a significant effect of expectations, with repetition enhancement being more pronounced for unexpected as compared to expected repetitions. Future studies with tailored paradigms optimized for detection of both repetition suppression and repetition enhancement could shed more light onto this unexpected finding.
